# Robust detection of clinically relevant features in single-cell RNA profiles of patient-matched fresh and formalin-fixed paraffin-embedded (FFPE) lung cancer tissue

**DOI:** 10.1101/2023.04.25.538273

**Authors:** Alexandra Trinks, Miha Milek, Dieter Beule, Julie Kluge, Stefan Florian, Christine Sers, David Horst, Markus Morkel, Philip Bischoff

## Abstract

Single-cell transcriptional profiling reveals cell heterogeneity and clinically relevant traits in intra-operatively collected patient-derived tissue. However, the established approach to perform such analyses on freshly collected tissue constitutes an important limitation since it requires prospective collection and immediate processing. Therefore, the ability to perform single-cell RNA sequencing from archived tissues would be very beneficial in a clinical setting. Here, we benchmark single-cell gene expression profiles from patient-matched fresh, cryopreserved and FFPE cancer tissue. We find that fresh tissue and FFPE routine blocks can be employed for the robust detection of clinically relevant traits on the single-cell level. Specifically, single-cell maps of fresh patient tissues and corresponding FFPE tissue blocks could be integrated into common low-dimensional representations, and cell subtype clusters showed highly correlated transcriptional strengths of signaling pathways, Hallmark and clinically useful signatures, despite some variability in expression of individual genes due to technological differences. FFPE tissue blocks revealed higher cell diversity compared to fresh tissue. In contrast, single-cell profiling of cryopreserved tissue was prone to artifacts in the clinical setting. Our analysis suggests that single-cell RNA sequencing from FFPE tissues is comparable to and can replace analyses from fresh tissue. This highlights the potential of single-cell profiling in the analysis of retrospectively and prospectively collected archival pathology cohorts and dramatically increases the applicability in translational projects.

## Introduction

High-throughput single-cell RNA sequencing (scRNA-seq) has enabled researchers to study the various aspects of cellular heterogeneity of tissues, including human clinical samples. In particular, scRNA-seq has been used extensively to characterize the complex cell compositions of tumor tissue cohorts ^1–3^, and can reveal features which contribute to patient prognosis ^4,5^ and therapy resistance ^6^. So far, single-cell RNA profiling required the prospective collection of fresh tissue samples during surgery or biopsy. Retrospective analysis of frozen or formalin-fixed and paraffin-embedded (FFPE) tissue cohorts would allow faster correlation of scRNA-seq profiles to clinical features, as many important clinical characteristics, such as tumor genetics, therapy response and patient survival, are only obtainable weeks to years after sample acquisition. In addition, this could unlock archival FFPE tissue collections for single-cell profiling.

Recently, scRNA-seq chemistry has been developed for frozen and formalin-fixed single-nucleus suspensions. Early studies have shown that this allows performing single-nucleus RNA sequencing (snRNA-seq) of nuclei isolated from frozen and FFPE tissue samples ^7–10^ at read depths that allow a similarly fine-grained analysis compared to fresh cell suspensions. However, to our knowledge, no side-by-side comparisons have been performed on fresh, frozen, and FFPE clinical tissue samples from the same patients. To answer the question which features of fresh tissue are preserved in archival samples, we profiled three lung adenocarcinomas using fresh, cryopreserved and FFPE tissue samples side-by-side. We found that cell-intrinsic and clinically relevant features of cancers are robustly preserved in single-cell transcriptomes of FFPE samples.

## Methods

### Sample acquisition

Tissue samples were collected from therapy-naïve lung adenocarcinoma patients undergoing primary surgery. Informed consent was obtained from all participants before sample acquisition. The ethics committee of Charité - Universitätsmedizin Berlin approved the use of tissue samples for single-cell gene expression analysis (vote EA4/164/19). Fresh tissue was sampled during intraoperative pathology consultation. Tissue samples were immediately snap-frozen in liquid nitrogen and stored at -80°C or placed in Tissue Storage Solution (Miltenyi) on ice for max. 30 min before subsequent further processing of the fresh tissue. The remaining surgery specimen was subjected to routine histological examination, including formalin fixation and paraffin embedding. FFPE tissue was sampled from archive tissue blocks after storage under standard archive conditions (dry, room temperature) for 4-5 months.

### Processing of fresh tissue samples

Fresh tissue samples were cut into small pieces of 1 mm diameter and dissociated using the Tumor Dissociation Kit, human (Miltenyi) and a gentleMACS Octo Dissociator with heaters (Miltenyi), using preinstalled program 37C_h_TDK_1 for 30-45 min. All subsequent steps were performed at 4 °C or on ice. Dissociated tissue was filtered through 100 µm filters, pelleted by centrifugation at 300 × g in BSA-coated low-binding tubes, incubated with 1 ml ACK erythrocyte lysis buffer (Gibco) for 1 min, washed with DMEM, pelleted, resuspended in PBS and filtered through 20 µm filters. Finally, debris was removed using the Debris Removal Solution (Miltenyi) and cells were counted using a Neubauer chamber. Single-cell suspensions were further processed using the Chromium Single Cell 3′ Reagent Kit v3 and the Chromium Controller (10x Genomics, Pleasanton, California, USA) according to the manufacturer’s protocol without any adjustments.

### Processing of cryopreserved tissue samples

Cryopreserved tissue samples were cracked using a pestle and mortar placed on dry ice and pre-chilled with liquid nitrogen into small pieces of 1 mm diameter. All subsequent steps were performed at 4 °C or on ice. Tissue pieces were homogenized in homogenization buffer (1x Nuclei EZ lysis buffer (Sigma), 0.6 U/ml RNAse Inhibitor (Ambion), 0.3 U/ml Superasin (Ambion) using a pestle and douncer by approx. 10 strokes with a loose pestle and 5 strokes with a tight pestle. Homogenized tissue was filtered through a 30 µm filter, stained with DAPI (0.1 µg/mL) for 5 min and sorted using a BD FACSAria Fusion (100 µm nozzle) to remove debris and doublets. Cell concentration was determined using a Neubauer chamber before single-nuclei suspensions were further processed using the Chromium Single Cell 3′ Reagent Kit v3 and the Chromium Controller iX (10x Genomics, Pleasanton, California, USA) according to the manufacturer’s protocol without any adjustments.

### Processing of FFPE tissue samples

FFPE tissue samples were processed according to a published protocol^8^ with adjustments according to a another published protocol^9^. In short, 4-6 tissue cores of 1 mm diameter were punched out of FFPE tissue blocks after the tumor area had been marked by a pathologist. Tissue cores were cut into small pieces of 1 mm diameter, washed at room temperature in xylene 3 times for 10 min, in 100 % ethanol 2 times for 30 sec, in 70 % ethanol for 30 sec, in 50 % ethanol for 30 sec, in distilled water for 30 sec, and in Buffer V (FFPE tissue dissociation kit, Miltenyi) for 30 sec. Tissue was dissociated using the FFPE tissue dissociation kit (Miltenyi) and a gentleMACS Octo Dissociator with heaters (Miltenyi), using preinstalled program 37C_FFPE_1. After 20 min of dissociation and after complete dissociation, dissociated tissue was pipetted through a 20G needle for 10-20 times. Next, samples were filtered through 70 µm filters, placed on ice, washed with chilled Buffer V, resuspended in Resuspension Buffer (0.5x PBS, 50mM Tris pH8, 0.02% BSA, 0.24U/µl RNasin (Ambion, AM2684) in H_2_O), and cells were counted using a Neubauer chamber. Single-nuclei suspensions were further processed using the Chromium Fixed RNA Profiling Reagent Kit and the Chromium Controller Xi (10x Genomics, Pleasanton, California, USA) according to the manufacturer’s protocol. After step 2.1.m in the manufacturer’s protocol, cells were stained with DAPI (0.1 µg/mL) for 5 min and sorted using a BD FACSAria Fusion (100 µm nozzle) to remove debris and doublets. After cell sorting, samples were further processed according to the manufacturer’s protocol.

### Library preparation and sequencing

After isolation, 10,000 cells/nuclei were subjected to barcoding and library preparation. Libraries of fresh and cryopreserved samples were prepared using the Chromium Single Cell 3′ Reagent Kit v3 (10x Genomics) according to the manufacturer’s protocol. Libraries of FFPE samples were prepared using the Chromium Fixed RNA Profiling Reagent Kit (10x Genomics) according to the manufacteurer’s protocol. Libraries were sequenced on a NovaSeq (Illumina) at approx. 400 mio. reads per library.

### Data analysis

Sequencing reads were aligned against reference transcriptome GRCh38 and UMIs were quantified using Cellranger, version 7.1.0 (10x Genomics). Subsequent analyses were performed using R version 4.1.1 and the Seurat package version 4.3.0 ^11^, if not stated otherwise. First, signal from ambient RNA was removed using the SoupX package version 1.6.2 ^12^, assuming a contamination fraction of up to 0.2. Gene expression data of all samples were merged and filtered for the following quality parameters: 300 to 10,000 genes per cell, 500 to 100,000 UMIs per cell, fraction of mitochondrial reads lower than 15 % and fraction of hemoglobin reads lower than 5 %. Next, gene expression data was log-normalized and dimensionality reduction was performed by principal component analysis (PCA). Uniform manifold approximation projection (UMAP) based on the top 10 principal components (PCs) was used for data visualization. Gene expression data was integrated by sample type (fresh, frozen or FFPE) using reciprocal PCA based on the top 30 PCs. Clustering was performed using shared nearest neighbor graph calculation. In fresh and FFPE data, main cell types and cell subtypes were manually annotated using canonical marker genes selected from the literature. Cell type labels were transferred based on the first 30 PCs to the frozen data. Clusters containing cell doublets were identified by discrepant marker gene expression and removed prior to further analysis. Signaling pathway activity scores were calculated using the PROGENy package version 1.17.3 ^13^. Gene signature expression scores were calculated using Seurat. The inferCNV package version 1.10.1 (https://github.com/broadinstitute/inferCNV) was used for copy number analysis in epithelial transcriptomes.

### Statistical analysis

The quantity of main cell types in fresh and FFPE samples were compared using the paired t test. The correlation of PROGENy pathway and Hallmark signature scores in different sample types (fresh, frozen, FFPE) was analyzed by calculating the Pearson correlation coefficient. Differentially expressed genes were identified using the FindAllMarkers function of the Seurat package with the following parameters: include only positive markers, proportion of expressing cells inside the cluster ≥ 0.2, difference between proportions of expressing cells inside and outside the cluster ≥ 0.2.

## Results

### Quality metrics of single-cell transcriptomes from fresh, frozen and archival FFPE samples

We selected cryopreserved and FFPE tissue samples of three lung adenocarcinoma patients (P075, P078, P079) that were previously analyzed by fresh tissue scRNA-seq for snRNA-seq analysis (Fig. 1A). Initial analysis of fresh and FFPE tissue-derived libraries showed expected high quality parameters (Fig. S1A); however, results gained from cryopreserved tissue showed unequal and lower quality scores in two subsequent rounds of library preparation and optimization of lab workflows (Fig. S1B). Across the patients, numbers of detected genes per sample were highest for fresh tissue, but lower for FFPE or frozen tissue (Fig. 1B), even when adjusting for the restricted gene set used in the targeted FFPE sequencing approach. Single-cell information of the fresh, frozen and FFPE samples could be integrated into a common UMAP with clusters sharing transcriptomes of fresh, frozen and FFPE tissue origin, indicating that cell type information was stable across the methods (Fig. 1C, Fig. S1C). When calling main cell types, we found that immune cell transcriptomes were enriched, but epithelial and stromal cells transcriptomes were rather depleted from fresh tissue single-cell libraries (Fig. 1D), indicating potential negative effects of tissue dissociation on cell representation. In line with this interpretation, a recently published gene signature for dissociation stress ^14^ was highest in fresh tissue transcriptomes, in particular in epithelial cells (Fig. 1E).

**Figure 1:**
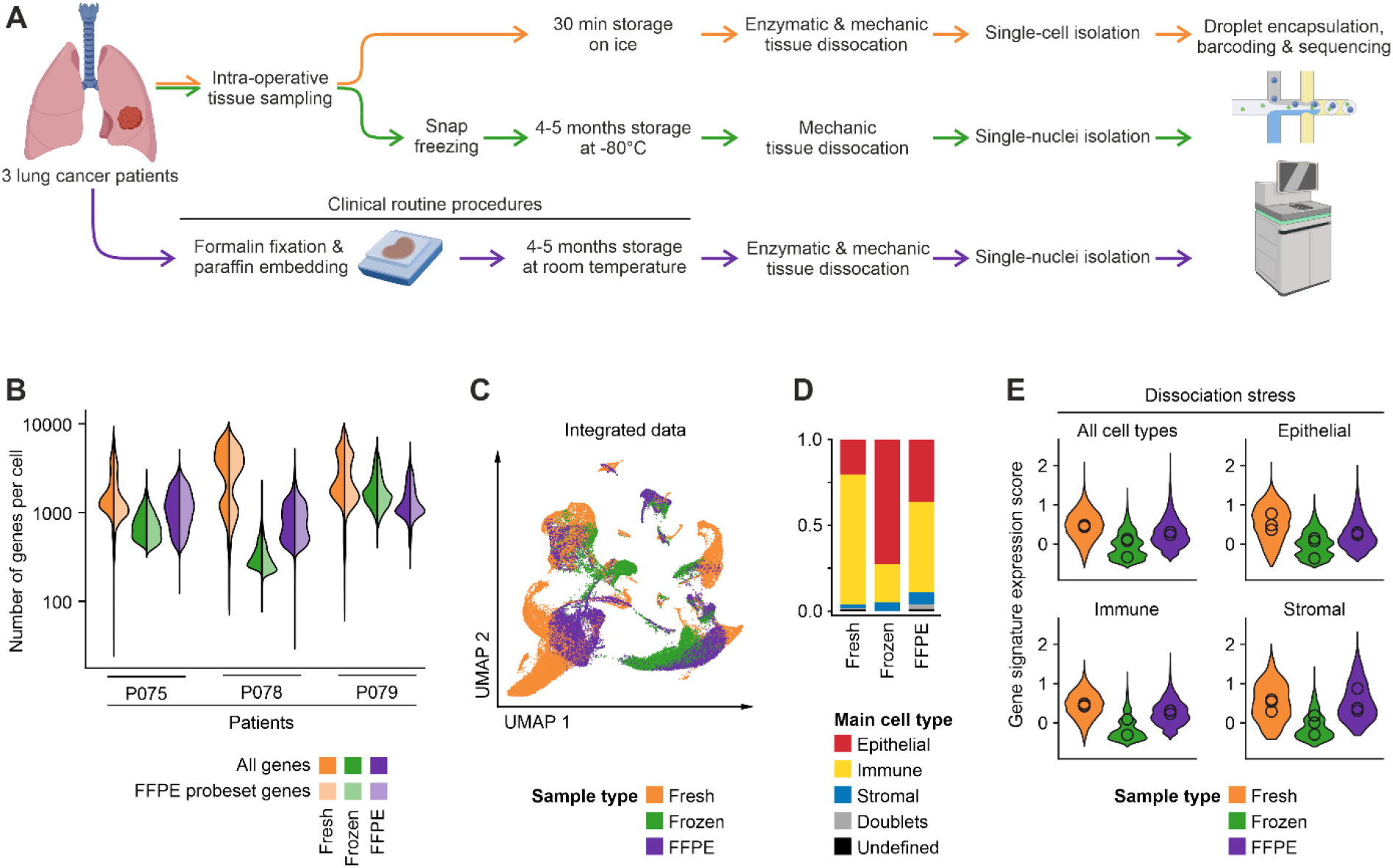
Workflow and quality metrics of fresh, frozen and FFPE tissue single-cell analysis. **A** Workflow of tissue specimens used for the study. In short, fresh tissues were procured intra-operatively, dissociated enzymatically, and cell suspensions were used for single-cell library preparation. Cryopreserved tissues were stored for 4-5 months at - 80°C, homogenized in the frozen state for nuclei isolation, and nucleus suspensions were used for library preparation. For FFPE analysis, 4-5 months old routine FFPE blocks were dissociated enzymatically and mechanically, and nucleus suspensions were used for library preparation. **B** Numbers of genes called per cell in the various libraries. Full colors: all genes; lighter colors: genes limited by FFPE probe set. **C** UMAPs based on the top 10 principal components of all single-cell transcriptomes after filtering and data integration, color-coded by fresh, frozen or FFPE tissue origin. **D** Quantification of main cell type, by clustering and calling of cell type-specific marker genes. **E** Module score of gene signatures related to dissociation stress in the various main cell types, by fresh, frozen and FFPE origin.

### Cell type abundancies in FFPE versus fresh samples

As FFPE tissue blocks are most relevant for clinical applications and frozen tissue libraries were hampered by low quality parameters, we integrated fresh and FFPE data for in-depth analysis (Fig. 2A), and again determined fractions of transcriptomes of epithelial, immune and stromal origin (Fig. 2B). Assessment of epithelial marker genes showed similar expression scores across fresh and FFPE tissue samples, as *EPCAM, KRT5, SCGB3A2, FOXJ1, AGER* and *SFTPC* marked tumor epithelial, Basal, Transitional, Ciliated, AT1, and AT2 cells, respectively, across the datasets (Fig. 2C). However, we also saw that some marker genes were preferentially detected in fresh versus FFPE transcriptomes, such as *MUC5B* and *SFTPC* in fresh or FFPE-derived tumor epithelial cells, respectively. This was likely due to the different absolute numbers of epithelial cell transcriptomes across fresh versus FFPE tissues (Fig. 2B), but also due to higher epithelial cell type diversity in the FFPE tissue blocks of patients P078 and P079 (Fig. 2D). The latter result suggests that normal epithelial cells are most easily lost during fresh tissue dissociation; however, we cannot rule out compositional bias in the fresh versus FFPE tissue specimens used. Detection of immune cell marker genes and cell types was more even across fresh and FFPE tissue specimens (Fig. 2E, F), while many more fibroblasts were detected among the stromal cell types in FFPE tissue-derived sequence libraries (Fig. 2G, H). Cell type label transfer to the cryopreserved tissue samples showed often skewed representation of cell types (Fig. S2A, B), in particular for immune cells such as regulatory or CD8+ T cells that are defined by few key marker genes which could not reliably detected in the frozen samples.

**Figure 2:**
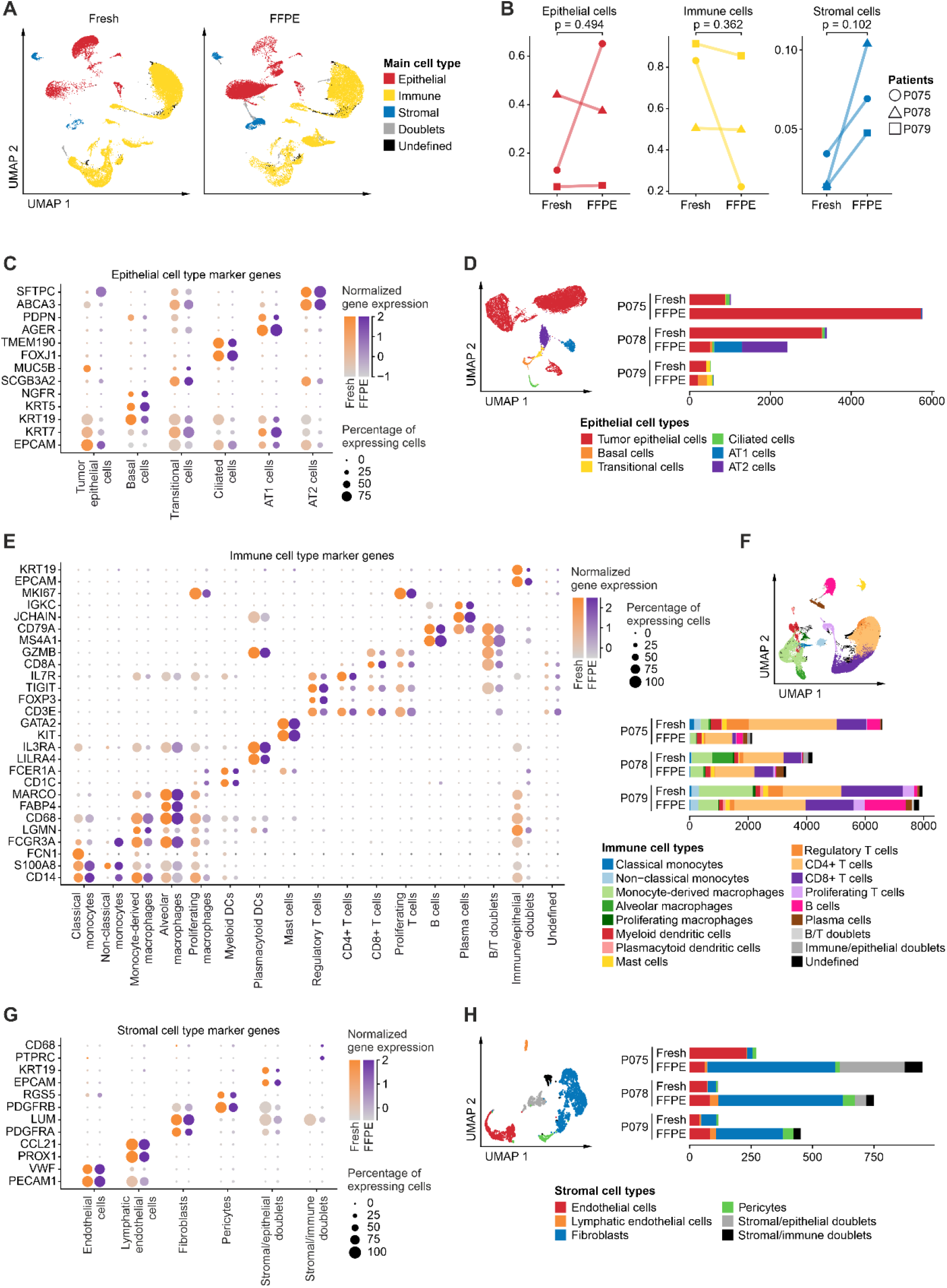
Cell type diversity in fresh versus FFPE tissue single-cell analysis. **A** UMAPs based on the top 10 principal components of fresh or FFPE single-cell transcriptomes, as indicated, color-coded by main cell type. **B** Relative proportions of epithelial, immune or stromal cells, compared between fresh and FFPE-derived libraries, paired t-test per main cell type. **C, D** Analysis of epithelial transcriptomes. **C** Epithelial marker gene expression per cell type in fresh or FFPE tissue-derived libraries. **D** UMAP and absolute cell numbers, color-coded by cell type. **E, F** Analysis of immune cell transcriptomes. **E** Immune marker gene expression per cell type in fresh or FFPE tissue-derived libraries. **F** UMAP and absolute cell numbers, color-coded by cell type. **G, H** Analysis of stromal cell transcriptomes. **G** Stromal marker gene expression per cell type in fresh or FFPE tissue-derived libraries. **H** UMAP and absolute cell numbers, color-coded by cell type.

### Cell trait heterogeneity in FFPE versus fresh samples

We asked whether archival FFPE tissue analysis allows characterization of cell types to a similar extent as fresh tissue single-cell analysis. We performed this analysis on gene signature level, taking into account sets of pathway target gene signatures ^13^ and relevant Hallmark gene sets ^15^. Technological differences in sample preparation workflows resulted in differences of normalized gene expression scores between the fresh tissue and the FFPE-derived libraries (Fig. S3A, B), where, for instance, the MAPK target gene *DUSP4* was found higher expressed in fresh tissue libraries, whereas *KMT5A* scored higher in FFPE-derived libraries. Despite these differences on individual gene expression levels, we found high correlations between inferred pathway activities (Fig. 3B) and tumor-related cell traits (Fig. 3C) from fresh and FFPE samples throughout. In line with prior studies, we found high TGFβ activity in fibroblasts, and this cell type also had the highest score for mesenchymal traits judged by activity of the EMT and Angiogenesis Hallmark signature. As expected, highest activity of the G2/M checkpoint and E2F targets Hallmark signatures was found in proliferating T cells and macrophages, which correlated with high MAPK activity in proliferating T cells. We detected high activity of TNFα and NFκB signaling in monocytes, and high scores of the Interferon Gamma Response Hallmark signature in monocytes and tumor-associated monocyte-derived macrophages (Mo-Macs), while tissue-resident alveolar macrophages were characterized by high scores of the Oxidative Phosphorylation Hallmark signature. In summary, correlations between signature activities derived from fresh or FFPE tissue samples were strong on the cell type level with Pearson correlation coefficients mostly larger than 0.9 and significances mostly smaller than p = 0.001 across all signatures analyzed, showing that biology-related cell type characteristics can be inferred from fresh and archival FFPE tissue specimens alike. Scores were lower when comparing correlations of cell type traits between fresh versus frozen libraries, in particular for pathways with smaller variation between cell types (Fig. S2C).

**Figure 3:**
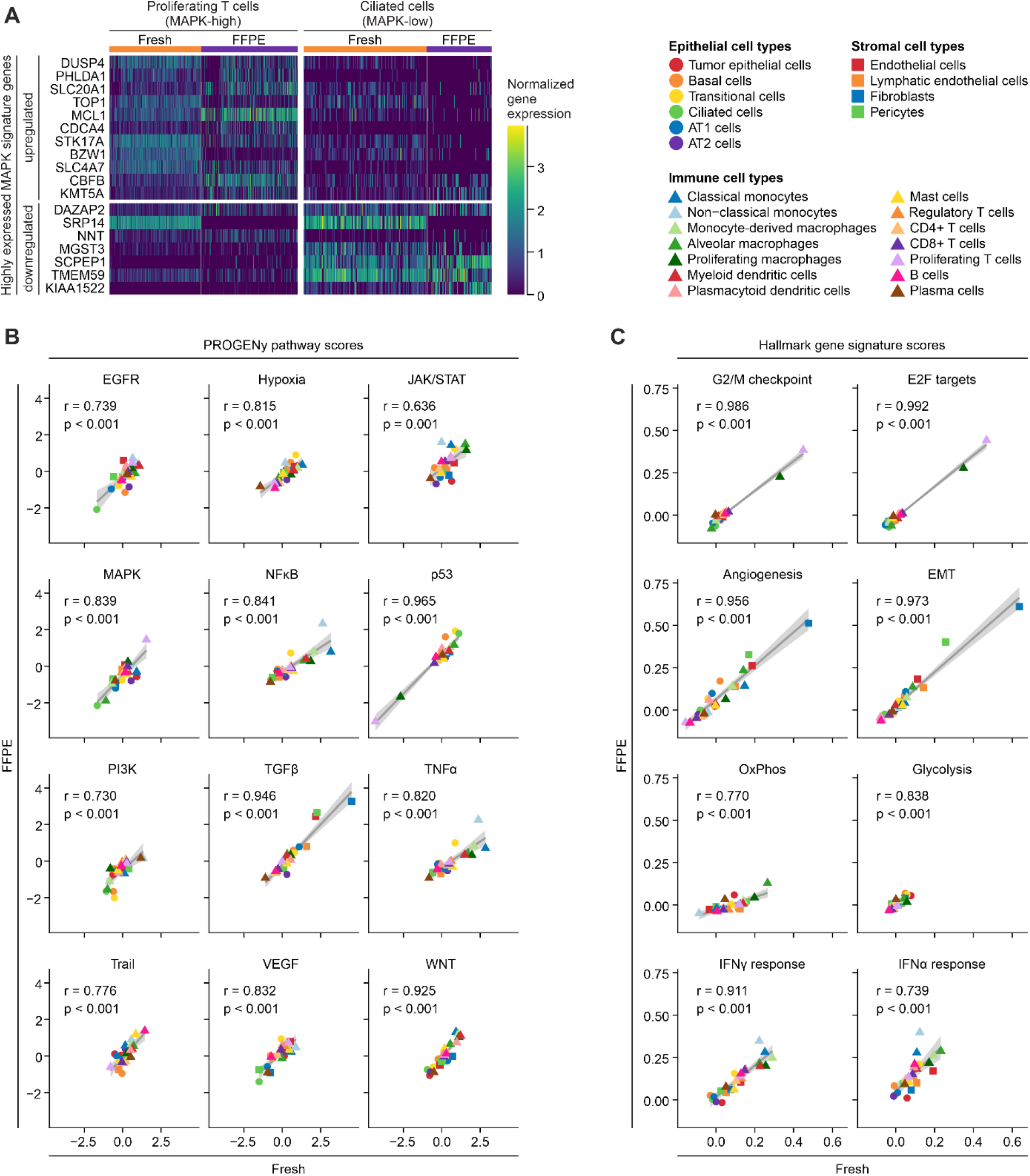
Cell trait quantification in fresh versus FFPE single-cell analysis. **A** Expression of selected PROGENy signature MAPK target genes in fresh tissue and FFPE libraries. Expression was normalized to scale across libraries. Data shown for cells assigned as Proliferating T cells (high MAPK activity, see Fig 3b) or Ciliated cells (low MAPK activity, see Fig 3b). **B** Correlations of PROGENy pathway scores between FFPE and fresh tissue gene expression per cell type, Pearson correlation coefficient and p-value indicated per pathway. **C** Correlations of selected biologically-relevant Hallmark gene signatures between FFPE and fresh tissue gene expression per cell type, Pearson correlation coefficient and p-value indicated per gene signature.

### Assessment of clinical features from FFPE versus fresh tissue single-cell transcriptomes

We investigated whether patient-specific differences in tumor and microenvironmental characteristics can be determined in fresh and FFPE tissue samples to the same degree. The histological grade of tumors (Fig. 4A) perfectly correlated with recently identified gene signatures of tumor differentiation ^16^ (Fig. 4B), regardless of whether the signatures were called from fresh or FFPE tissue-derived epithelial cell transcriptomes. The lowest histological tumor grade in patient P079 was further associated with higher activity of CAF-related pathways in fibroblasts^17^ (Fig. 4C) and high expression of typical CAF marker genes on the single-cell mRNA level (Fig. 4D). With regard to oncogenic pathways, tumor epithelial cells of patient P079 scored highest for MAPK and WNT target expression, while P075 had the highest values of p53 pathway activity, in both fresh and FFPE tissue samples (Fig. S4A). In contrast, no correlation was observed for EGFR pathway activity. We inferred similar copy number profiles in tumor epithelial cells from both fresh and FFPE tissue-derived transcriptomes (Fig. S4B).

**Figure 4:**
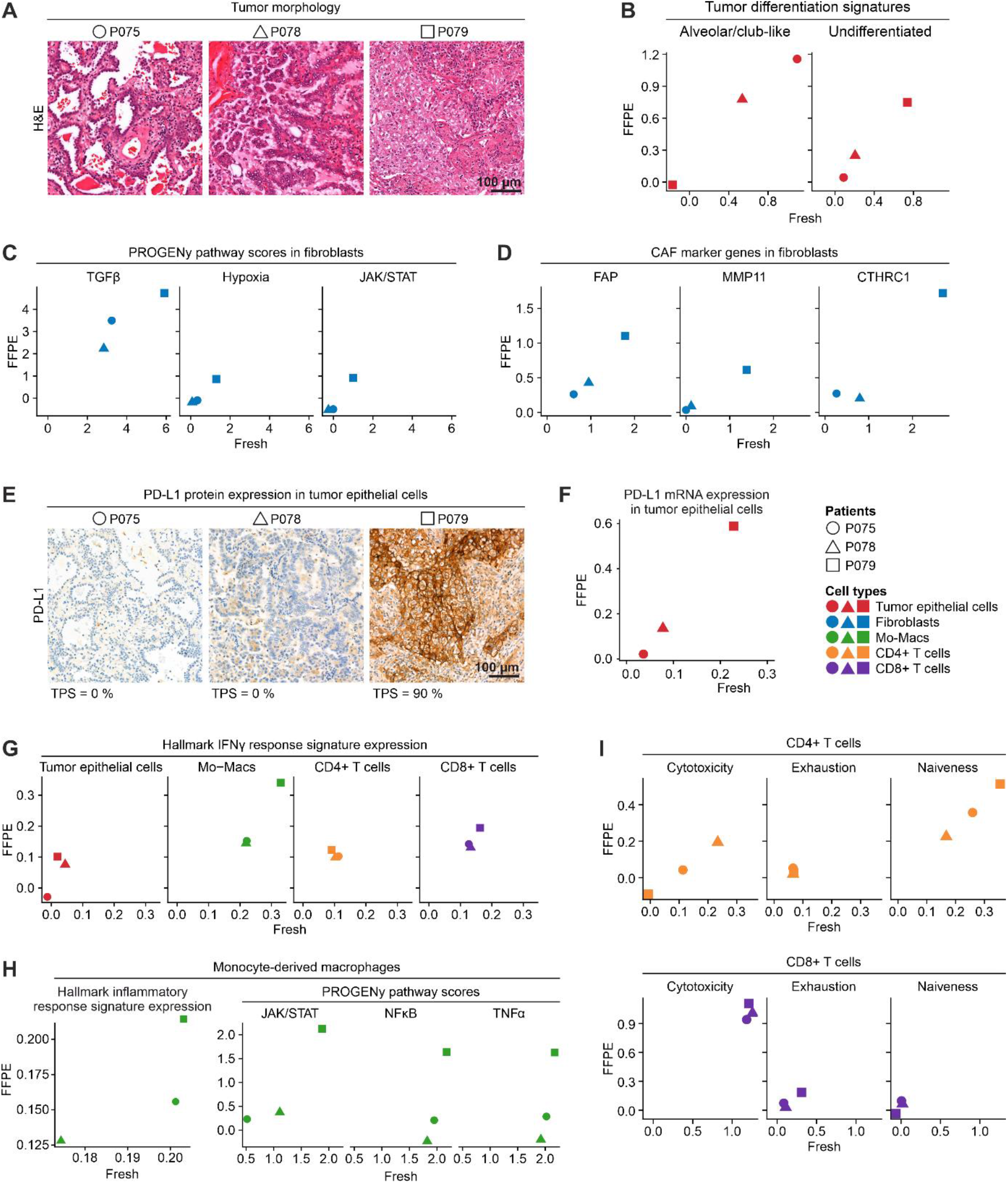
Quantification of clinically relevant gene expression patterns in fresh versus FFPE single-cell analysis. **A** Hematoxylin and eosin stained FFPE tumor sections of patients P075, P078, P079. **B** Correlation between FFPE and fresh tissue of Alveolar/club-like and Undifferentiated tumor cell signature expression in tumor epithelial cells. **C** Correlation between FFPE and fresh tissue of PROGENy pathway scores in fibroblasts. **D** Correlation between FFPE and fresh tissue of CAF marker expression in fibroblasts. **E** Immunohistochemistry on FFPE tumor section for PD-L1 expression. TPS: Tumor proportion score. **F** Correlation between FFPE and fresh tissue of PD-L1 gene expression in tumor epithelial cells. **G** Correlation between FFPE and fresh tissue of Interferon Gamma Response Hallmark gene signature expression in various cell types, as indicated. Mo-Macs = Monocyte-derived macrophages. **H** Correlation between FFPE and fresh tissue of clinically and biologically relevant signature scores in Mo-Macs, as indicated **I** Correlation between FFPE and fresh tissue of functional cell state scores in CD4+ and CD8+ T cells, as indicated.

Histologically, tumors differed in expression of the immune evasion marker PD-L1 in tumor cells (Fig. 4E) which was similarly observed on the RNA level in fresh as well as FFPE tissue-derived tumor epithelial cell transcriptomes (Fig. 4F). In both fresh and FFPE transcriptome data, high PD-L1 expression in patient P079 correlated with high expression of the Interferon Gamma Response Hallmark signature in tumor cells and the tumor microenvironment, in particular in Mo-Macs^18^ (Fig. 4G). In Mo-Macs, this feature was accompanied by high TNFα, NFκB and JAK/STAT pathway activity and high expression scores of the Inflammatory Response Hallmark signature in patient P079 (Fig. 4H)^19,20^. Among T cells, high PD-L1 expression in P079 correlated with low cytotoxicity and high naiveness scores in CD4+ T cells, and high scores for exhaustion in CD8+ T cells (Fig. 4I)^21^. In summary, these correlations are in agreement with previous analysis of the lung cancer microenvironment ^16,22^, and indicate that cell type characteristics relevant for clinical stratification can be retrieved faithfully from the FFPE snRNA-seq approach.

## Discussion

Single-cell sequencing of surgical tissues has uncovered clinically relevant information on the patient and cohort levels. However, the use of fresh tissues comes with serious limitations in the clinic, such as the requirement for a rapid tissue handling pipeline. Recent technological advances allow single-nucleus sequencing from frozen and FFPE tissue, which promises broader availability of tissue, including routine pathology specimens. Here, we present a systematic comparison of fresh, frozen and FFPE single-cell analysis of clinical tissue. We find that FFPE tissue robustly preserved clinically relevant information on cell types and patient characteristics comparable to fresh tissue.

Single-cell analysis of fresh solid tissue has been the gold standard to survey cell type composition of human tissues in health and disease ^23–25^. Generally, the procedure relies on cell dissociation, most often by proteases, which can result in disproportionate release of cell types and preparation-related transcriptome artifacts, although procedures have been developed to minimize such effects. Moreover, fresh tissue requires timely processing, which can complicate standardization and result in batch effects. Potentially, analysis of transcriptionally inert nuclei from frozen or FFPE tissues could result in more even transcriptome representation across cell types in the absence of dissociation artifacts. We have here tested available procedures side-by-side on tissues of the same origin and found that single-cell sequencing of FFPE tissue results in transcriptomes representing a higher diversity of cell types compared to analysis of fresh tissue cell suspensions, an equally good representation of biological and clinical transcriptome features on a signature level, and lower induction of stress-related gene expression. In contrast, frozen tissue analysis was less reliable in our hands, probably due to handling of unfixed nuclei suspensions that are prone to RNA degradation or RNA diffusion.

In the clinic, fresh tissue single-cell analysis allowed for prospective collection of samples only. Importantly, single nucleus transcriptomics of FFPE tissues allow for retrospective analysis of cohorts that are annotated with long-term clinical follow-up data such as therapy response, relapse, metastasis, and patient survival. The FFPE blocks under analysis here were stored under standard archival conditions at room temperature for 4-5 months. It remains to be determined whether older FFPE blocks with longer follow-up periods perform equally well, as RNA quality could potentially deteriorate over time even in the fixed and embedded state. Moreover, it needs to be investigated to which degree prolonged time until fixation and tissue autolysis in large surgery specimens impair snRNA-seq analysis.

Pathology review is usually performed using FFPE sections, as these yield high-quality histology and immunohistochemistry information. Therefore, FFPE blocks can be annotated regionally with high confidence, unlike intra-operative fresh tissue samples. Thus, FFPE blocks allow analysis of cell composition of circumscribed areas of interest. We envision that careful selection of tissue regions in pathology, in conjunction with multiplexing techniques for cost-effectiveness, will bring single-cell analysis aiding therapy prediction or prognosis a step closer to clinical routine for cancer patients.

## Data availability

Single-cell gene expression data generated in this study and analysis scripts are available on Zenodo at https://doi.org/10.5281/zenodo.7852154.

## Author Contributions

Conceptualization: PB, SF, MM; Investigation: AT, PB, JK; Data Curation: MiM, DB, PB; Formal Analysis: PB; Validation: PB; Visualization: PB, MM; Writing – Original Draft: PB, MM, AT; Resources: MM, PB, CS, DH

## Acknowledgments

Figure 1A was created with BioRender.com.

