## Supplemental Figures for "Robust detection of clinically relevant features in single-cell RNA profiles of patient-matched fresh and formalin-fixed paraffin-embedded (FFPE) lung cancer tissue"

### Supplementary figures

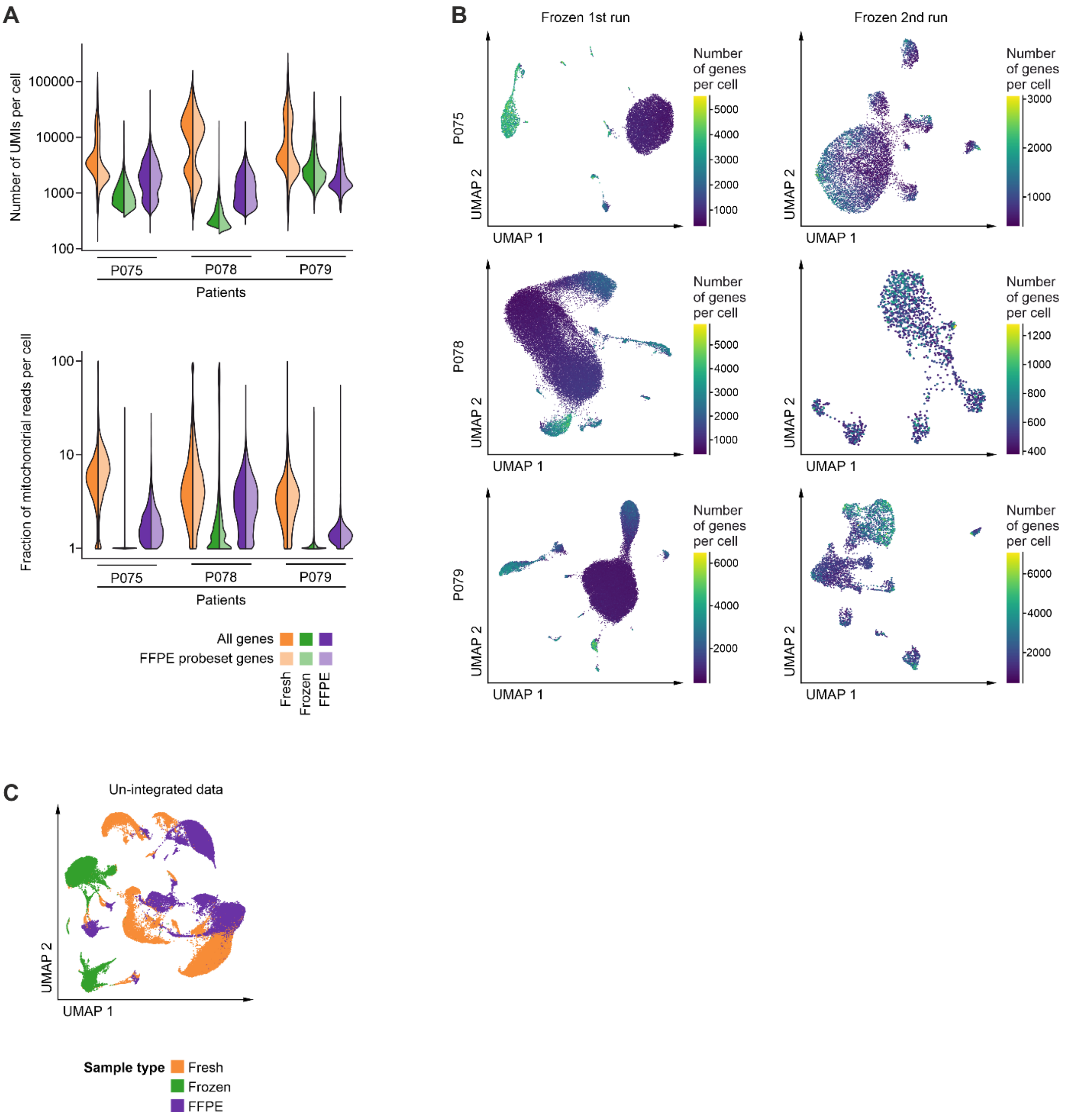


**Figure S1: Additional quality metrics of fresh, frozen and FFPE tissue. A** Numbers of unique molecule identifiers (UMIs) and fractions of mitochondrial reads per cell across libraries. Full colors: all genes; lighter colors: genes limited by FFPE probe set. **B** UMAPs of frozen tissue libraries showing insufficient cluster separation. Only patient P079, 2nd run shows sufficient cluster separation, and thus cell type annotation. **C** UMAPs based on the top 10 principal components of all single-cell transcriptomes after filtering and normalization, without data integration, color-coded by fresh, frozen or FFPE tissue origin. For UMAP of integrated data, see main Fig. 1C.

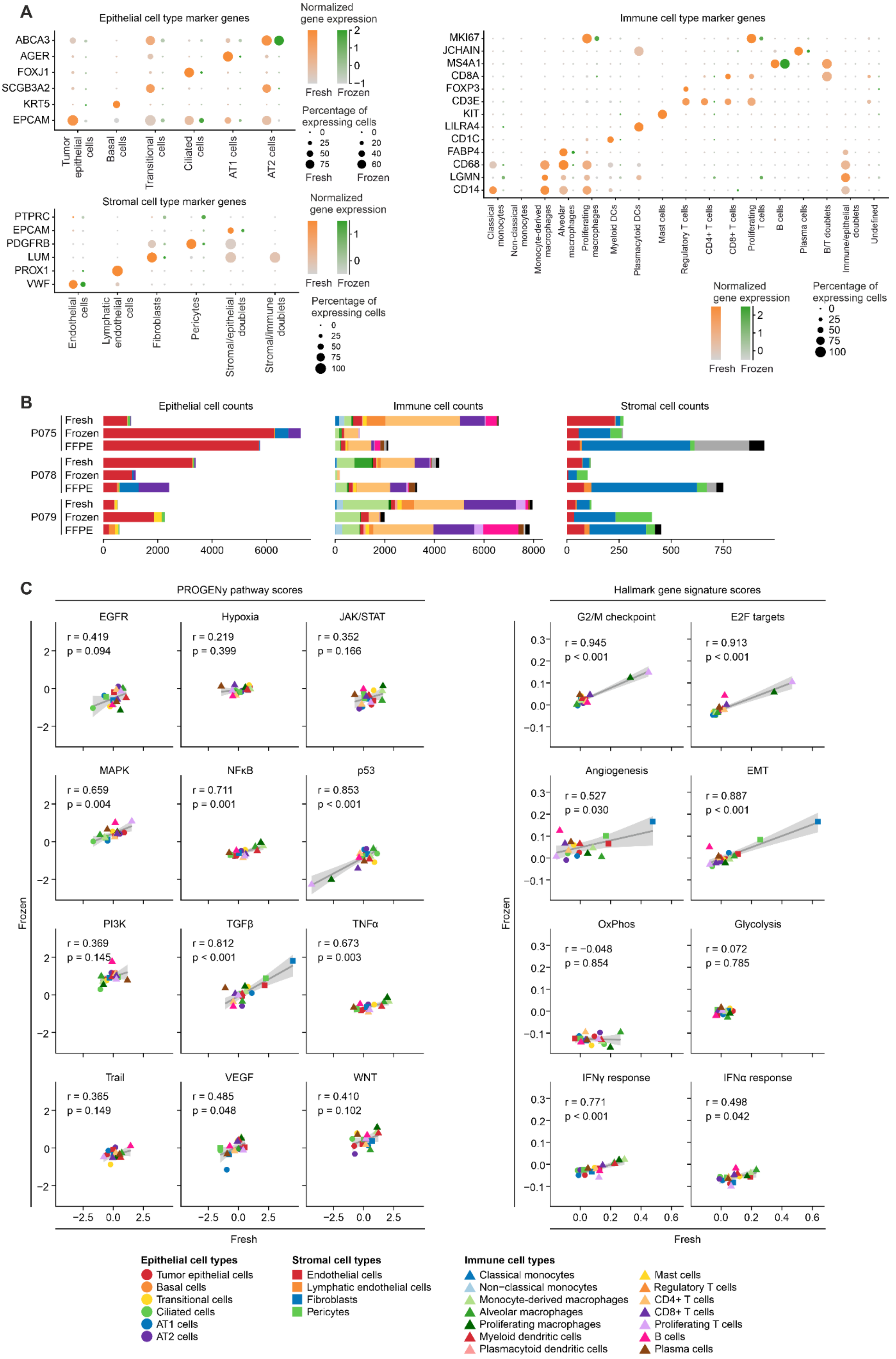
A
Epithelial cell type marker genes
ABCA3
AGER
FOXJ1
SCGB3A2
KRT5
EPCAM
Tumor epithelial cells
Basal cells
Transitional cells
Ciliated cells
AT1 cells
AT2 cells
Normalized gene expression
Fresh Frozen
Percentage of expressing cells
Stromal cell type marker genes
PTPRC
EPCAM
PDGFRB
LUM
PROX1
VWF
Endothelial cells
Lymphatic endothelial cells
Fibroblasts
Pericytes
Stromal/epithelial doublets
Stromal/immune doublets
Immune cell type marker genes
MKI67
JCHAIN
MS4A1
CD8A
FOXP3
CD3E
KIT
LILRA4
CD1C
FABP4
CD68
LGMN
CD14
Classical monocytes
Non-classical monocytes
Monocyte-derived macrophages
Alveolar macrophages
Proliferating macrophages
Myeloid DCs
Plasmacytoid DCs
Mast cells
Regulatory T cells
CD4+ T cells
CD8+ T cells
Proliferating T cells
B cells
Plasma cells
B/T doublets
Immune/epithelial doublets
Undefined
B
Epithelial cell counts
Immune cell counts
Stromal cell counts
P075 Fresh Frozen FFPE
P078 Fresh Frozen FFPE
P079 Fresh Frozen FFPE
C
PROGENy pathway scores
EGFR r = 0.419 p = 0.094
Hypoxia r = 0.219 p = 0.399
JAK/STAT r = 0.352 p = 0.166
MAPK r = 0.659 p = 0.004
NFκB r = 0.711 p = 0.001
p53 r = 0.853 p < 0.001
PI3K r = 0.369 p = 0.145
TGFβ r = 0.812 p < 0.001
TNFα r = 0.673 p = 0.003
Trail r = 0.365 p = 0.149
VEGF r = 0.485 p = 0.048
WNT r = 0.410 p = 0.102
Hallmark gene signature scores
G2/M checkpoint r = 0.945 p < 0.001
E2F targets r = 0.913 p < 0.001
Angiogenesis r = 0.527 p = 0.030
EMT r = 0.887 p < 0.001
OxPhos r = −0.048 p = 0.854
Glycolysis r = 0.072 p = 0.785
IFNγ response r = 0.771 p < 0.001
IFNα response r = 0.498 p = 0.042
Frozen
Fresh
Epithelial cell types
Tumor epithelial cells
Basal cells
Transitional cells
Ciliated cells
AT1 cells
AT2 cells
Stromal cell types
Endothelial cells
Lymphatic endothelial cells
Fibroblasts
Pericytes
Immune cell types
Classical monocytes
Non-classical monocytes
Monocyte-derived macrophages
Alveolar macrophages
Proliferating macrophages
Myeloid dendritic cells
Plasmacytoid dendritic cells
Mast cells
Regulatory T cells
CD4+ T cells
CD8+ T cells
Proliferating T cells
B cells
Plasma cells

**Figure S2: Cell type diversity in fresh, frozen and FFPE tissue single-cell analysis. A** Cell type marker gene expression per cell type in fresh or frozen tissue-derived libraries. **B** Absolute cell numbers per cell type, in fresh, frozen or FFPE-derived libraries. **C** Cell trait quantification in fresh versus frozen single-cell analysis. Correlations of PROGENy pathway or Hallmark signature scores are given between frozen and fresh tissue gene expression per cell type. Pearson correlation coefficient and p-value indicated per pathway or gene signature.

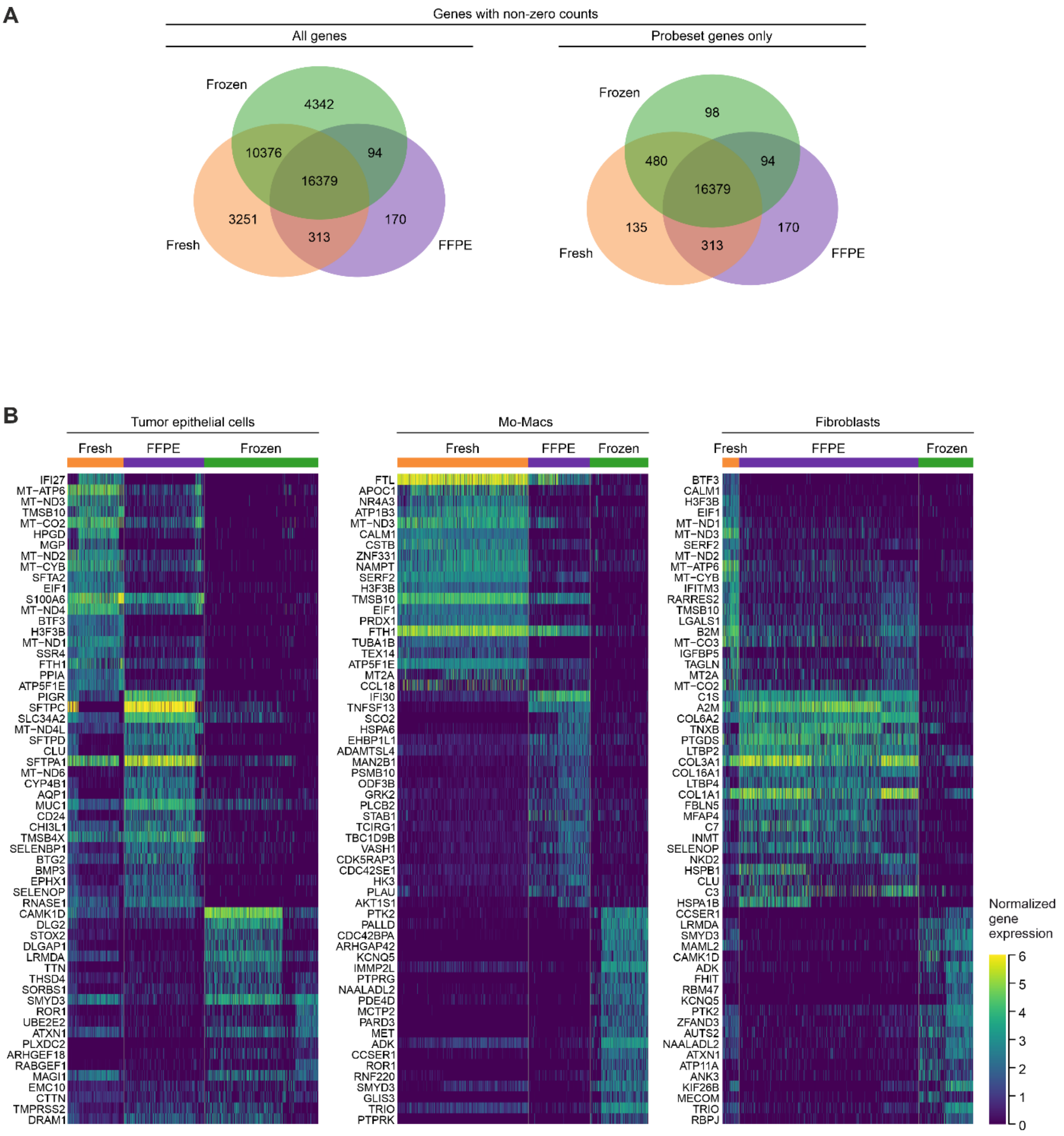


**Figure S3: Gene expression quantification across single-cell libraries. A** Numbers of genes detected (non-zero counts) across the technologies. To the left: all genes, to the right: genes represented by FFPE probe sets only. **B** Top 20 differentially expressed genes across the technologies in tumor epithelial cells, Monocyte-derived macrophages (Mo-Macs) and fibroblasts, respectively.

A

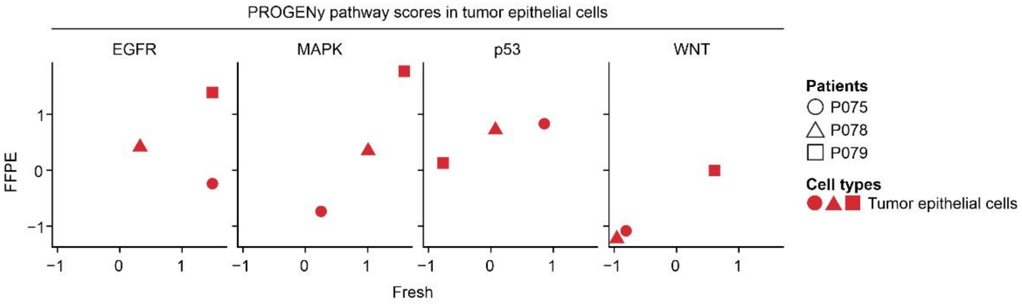
PROGENy pathway scores in tumor epithelial cells
EGFR
MAPK
p53
WNT
FFPE
Fresh
Patients
P075
P078
P079
Cell types
Tumor epithelial cells

B

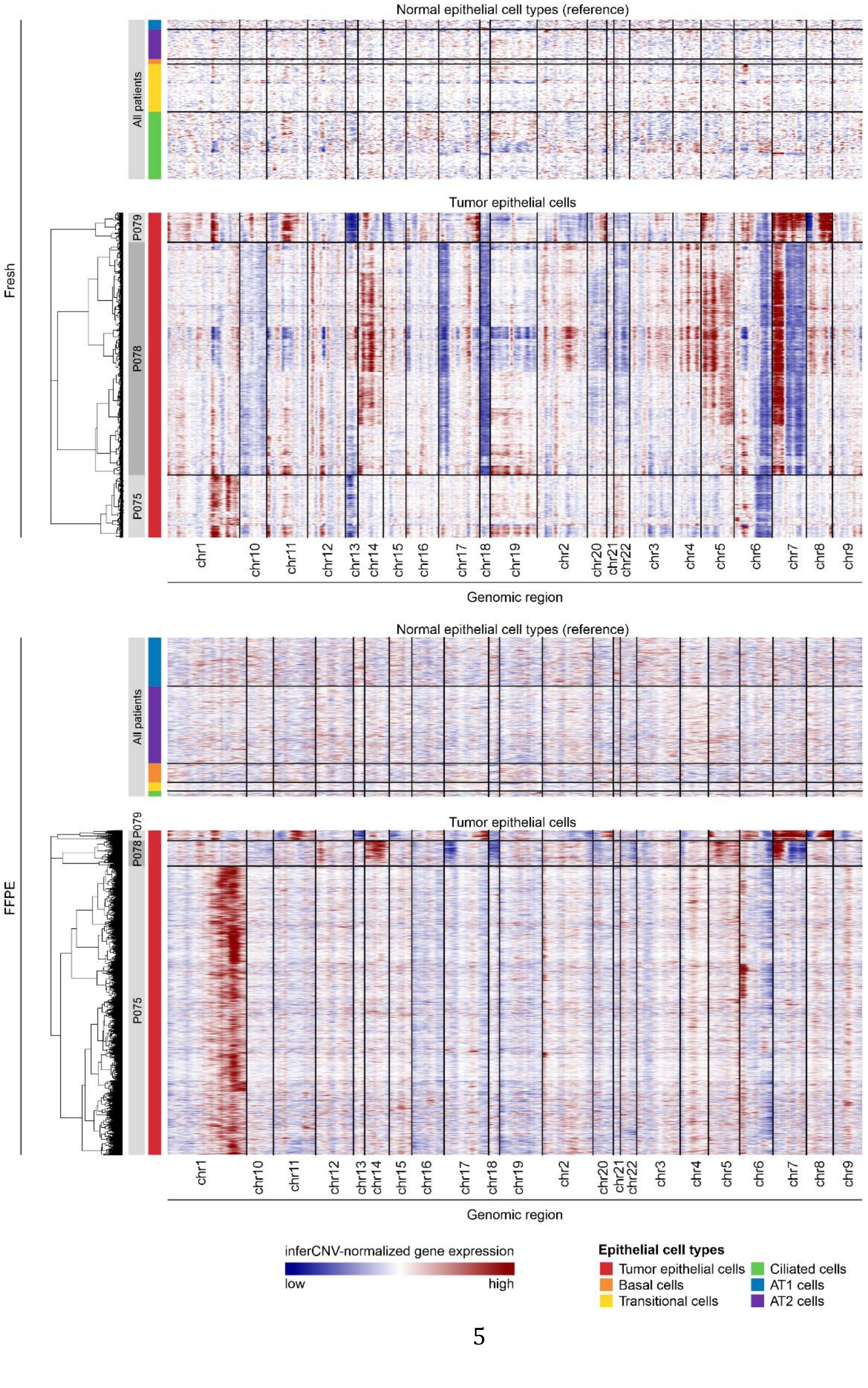
Normal epithelial cell types (reference)
All patients
Tumor epithelial cells
Fresh
P079
P078
P075
Genomic region
FFPE
inferCNV-normalized gene expression
low
high
Epithelial cell types
Tumor epithelial cells
Basal cells
Transitional cells
Ciliated cells
AT1 cells
AT2 cells

**Figure S4: Quantification of oncogenic pathway activities and inference of copy number aberrations in fresh versus FFPE single-cell analysis of tumor epithelial cells. A** Correlation between FFPE and fresh tissue of PROGENy EGFR, MAPK, p53 and Wnt target gene signatures, as indicated. **B** Predicted copy number aberrations inferred from tumor epithelial single-cell transcriptomes of fresh or FFPE tissue origin, as indicated.
